# Smartphone image capture system and image analysis pipelines enable accurate and efficient phenotyping of spaced plant mapping populations

**DOI:** 10.64898/2026.01.06.697986

**Authors:** Stella Woeltjen, Molly Hanlon, Keely Brown, Haley Schuhl, Ivan Baxter, Allison Miller

## Abstract

The lack of low-cost, user-friendly and expedient methods for plant phenotyping challenges researchers’ ability to efficiently collect accurate phenotypic data in large field experiments. Here, we demonstrate the use of a novel, smartphone-based image capture system and two user-friendly image analysis pipelines (utilizing PlantCV or Biodock AI) to increase the throughput of plant phenotyping in two large, spaced plant populations. We showed that the image capture system collected images of adequate quality for downstream analysis using either the PlantCV or Biodock AI pipeline. Both image analysis pipelines produced phenotype values in line with those obtained using manual image annotation. Together, these results provide researchers with a low-cost, user-friendly image-based phenotyping method that can be widely applied to increase phenotyping throughout in field experiments.

## Abstract

Adoption of in-field image-based plant phenotyping is hampered by a lack of low-cost, user-friendly image collection and analysis pipelines. In this study, we tested the utility of a low-cost, smartphone-based image capture system (SICS) for capturing overhead images of two herbaceous perennial plant species (*Silphium* L. backcross population and *Trifolium ambiguum* M. Bieb.) in field conditions. Two image analysis pipelines were developed using readily accessible image analysis softwares: PlantCV or Biodock AI. The ability of each pipeline to generate accurate estimates of plant area and plant area trends, while maintaining low image processing times, was evaluated. Using over 12,000 images collected over two field seasons, this study demonstrated the suitability of the SICS to capture images for plant trait analysis using either PlantCV or Biodock AI pipelines. Both pipelines enabled rapid extraction (minutes to hours) of plant trait information from images while maintaining high accuracy, particularly for simple image compositions (where a single plant was growing against a dark soil background), with plant trait estimates within 30% of the values generated by manual image analysis using ImageJ. Though accuracy declined marginally when analyzing complex images (where a single plant was growing with other plants in the background), the decline in accuracy did not significantly alter trends in mean plant area estimated by either pipeline. The methodological advancements presented in this study are expected to better support the widespread adoption of image-based plant phenotyping in field studies, allowing for the rapid advancement of germplasm in plant research and breeding programs.

## 1 Introduction

Plant phenotyping remains a bottleneck for plant research and breeding programs (Montes et al., 2007)., due, in part, to constraints posed by traditional phenotypic data collection methods (Araus et al., 2018; Araus and Cairns, 2014). Traditional manual measurement of plant traits in-field can be time consuming and cost-prohibitive. Further, manual plant phenotyping is prone to human error (Heineck et al., 2019) that can hamper the detection of phenotypic trends in plant populations (Araus et al., 2018). The situation is further complicated in perennial systems, where consistency in repeated measurements on the same plants across seasons and over years can be challenging. Alternative approaches to manual phenotyping, such as image-based phenotyping, may relieve the plant phenotyping bottleneck in experimental research and breeding programs (Song et al., 2021). Such methods can replace the need for manual phenotyping, increase both the efficiency and accuracy of plant phenotype collection, provide more consistent data collection, and alleviate the bottleneck created by manual phenotyping methods.

Despite the potential for image-based phenotyping to rapidly and accurately advance our biological knowledge in a variety of crops, current imaging systems are largely cost prohibitive, have a steep learning curve, or have been designed to work at scales not relevant or useful to answering questions in all systems (Hoyos-Villegas et al., 2025). For example, aerial image capture systems (i.e., satellites, uncrewed aerial vehicles (UAVs), etc.), are of broad interest for use in field image-based phenotyping due to the ability to rapidly collect imagery of large plot areas, but can be costly, are subject to federal and local regulations, and require specialized expertise to operate (Guo et al., 2021; Gano et al., 2024). Ground-sensing vehicles (GSVs) present a lower-barrier alternative to aerial image capture systems, with GSVs ranging in complexity from RGB cameras mounted to tractors to pull-behind imaging boxes attached to all-terrain vehicles (Kicherer et al., 2017; Sharma and Ritchie, 2015; Jiang et al., 2018; Busemeyer et al., 2013). However, similar to aerial image capture systems, the construction and operation of GSVs can incur prohibitive costs and still require specialized expertise and equipment that may not be accessible to all researchers. Deviation from a row-crop designed experiment can further complicate the use of UAVs or GSVs as fields may be more difficult to traverse or analyze in perennial, multi- or inter-cropped systems. These barriers limit the widespread adoption of image-based plant phenotyping in field studies and underscore the need for increased access to low-cost, low-barrier image capture systems.

Beyond image collection, the lack of readily accessible and expedient analysis pipelines to extract plant trait information from images further hampers the adoption of image-based phenotyping in the field (Minervini et al., 2015; Gano et al., 2024; Guo et al., 2021). The complexity of computational pipelines used to extract plant phenotypic information from images can take weeks to months to develop, prolonging the time between image capture and data availability (Gano et al., 2024). Further, many existing image capture systems and analysis pipelines are tailored to specific crops and capture methods with limited transferability to alternative settings. Complex image compositions, such as those containing non-target plants touching target plants, shadows in plant canopies, soil occluding leaf material, or some combination of these factors, often pose challenges to the automated or semi-automated extraction of plant trait information from images. As complex image compositions are common to most field settings, a paired image capture and analysis pipeline robust to complex field image conditions is needed to reduce turnaround times between image collection and data availability and promote adoption of field image-based phenotyping methods.

Recent advances in machine-learning based analysis methods have resulted in the development of new tools to expedite image analysis, but these tools are often not accessible to researchers lacking a background in computer vision or access to high-performance computing resources. PlantCV (Plant ComputerVision) and Biodock AI may help researchers overcome this hurdle by providing accessible image analysis software that allows for plant trait extraction from smartphone images, potentially balancing both ease-of-use with low turnaround time to plant trait data availability. PlantCV is an open-source, python-based software that enables automated extraction of traits from plant images (Gehan et al., 2017). At the time of publication, built-in PlantCV functions enable the extraction of over 100 image features, allowing for quantification of plant traits such as branching architecture, tissue color, and plant area. PlantCV allows for custom functions to be developed, enabling broad flexibility to meet the needs of a variety of researchers. PlantCV has been widely used to extract plant trait information from overhead and side-view images of whole plants grown under controlled conditions, including maize (Duenwald et al., 2025; Enders et al., 2019), Setaria (Feldman et al., 2018; Fahlgren et al., 2015), and Arabidopsis. A second image analysis tool is Biodock AI (Biodock, 2024), a semi-automated and non-programmatic option for plant trait extraction from images. Biodock AI is a web-based image analysis platform that allows users to train custom AI models (built on the Segment Anything Model (SAM)) that can be deployed across image sets to isolate target plants for trait analysis. Biodock AI has been used for image analysis applications in cellular research (Walter et al., 2024; Vijayakumar et al., 2023; Pannoni et al., 2025) and materials science (Alina et al., 2025). However, as PlantCV and Biodock AI were primarily developed to quantify features in images collected under conditions where image composition and lighting are highly controlled, it remains unclear whether PlantCV and Biodock AI image analysis pipelines are suitable for extracting plant trait information from images collected under field conditions.

With the goal of enhancing the efficiency and accuracy of phenotyping in plant research and breeding programs, in this study we aimed to evaluate two imaging pipelines that can be used to overcome image collection and analysis barriers to image-based phenotyping in the field The objectives of this study were to: 1) build a lightweight, simple, smartphone-based field image collection system; 2) use PlantCV and Biodock AI to develop and test two image analysis pipelines to extract plant trait information from images collected under field conditions; and 3) evaluate the accuracy of plant traits estimated using the PlantCV and Biodock AI image analysis pipeline to aid researchers in adopting such methods in plant research and breeding programs.

## 2 Materials and Methods

### 2.1 Plant Material and Field Layout

This study focused on repeated imaging in the field of individual plants from two perennial, herbaceous species: *Trifolium ambiguum* (M. Bieb) and an interspecific cross between *Silphium perfoliatum* x *Silphium integrifolium* (hereafter, ‘*Silphium’*). One *T. ambiguum* and one *Silphium* nursery were established at the Danforth Plant Science Center Field Research Site (St. Charles, MO) with individual plants spaced on a 76 cm grid (Table 1). The *T. ambiguum* nursery was established in fall 2022 and contains 442 genotypes replicated across four blocks, with a total of 1,768 individual plants. The *Silphium* nursery was established in spring 2023 and contains 700 individual plants. A second *Silphium* nursery was established in spring 2023 at the Shaw Nature Reserve (Gray Summit, MO) and includes 440 individual plants.

**Table 1.** Spaced plant population information, number of images collected to date, and imaging frequency and duration for each perennial plant species.

|  | *Silphium* | *Trifolium ambiguum* |
| --- | --- | --- |
| ***Population*** |  |  |
| Established | Spring 2023 | Fall 2022 |
| No. of Individuals | 1,135 | 1,768 |
| ***Images Collected*** |  |  |
| 2023 | 2,230 | 1,768 |
| 2024 | 1,115 | 5,304 |
| 2025 | 5,995 | 3,968 |
| ***Imaging Duration*** |  |  |
| Frequency | Weekly | Monthly |
| Growth Stage | Rosette / Vegetative | Vegetative to Flowering |
| Image Collection time | 6 min 100 plant<sup>-1</sup> | 24 min 100 plant<sup>-1</sup> |
| ***Image Pipeline Analysis Time*** |  |  |
| ImageJ Pipeline | 137 min 100 plant<sup>-1</sup> | 109 min 100 plant<sup>-1</sup> |
| PlantCV Pipeline | 168 min 100 plant<sup>-1</sup> | 243 min 100 plant<sup>-1</sup> |
| Biodock AI Pipeline | 21 min 100 plant<sup>-1</sup> | 18 min 100 plant<sup>-1</sup> |

### 2.2 Image Capture with Smartphone Image Capture System (SICS)

Overhead images of individual plants in the field were collected over three growing seasons using the Smartphone Image Capture System (SICS). The SICS is a custom-designed, lightweight, and inexpensive field image capture system composed of a smartphone (iPhone 14 Pro), a custom PVC color card holder, a color card (4 x 6 Pixel Perfect Standard Color Calibration Card), and a Bluetooth-enabled remote shutter (Figure 1). To capture images using the SICS, the smartphone was held directly above the target plant t from which traits were to be extracted. The smartphone was positioned so the target plant was centered in the image frame, and the color card was fully visible (Figure 2). Both the smartphone and color card were positioned parallel to the ground surface, and image capture was triggered using the remote shutter. Over the imaging period, a total of 11,040 and 9,340 images were collected from *T. ambiguum* and *Silphium* nurseries, respectively (Table 1).

**Figure 1.**
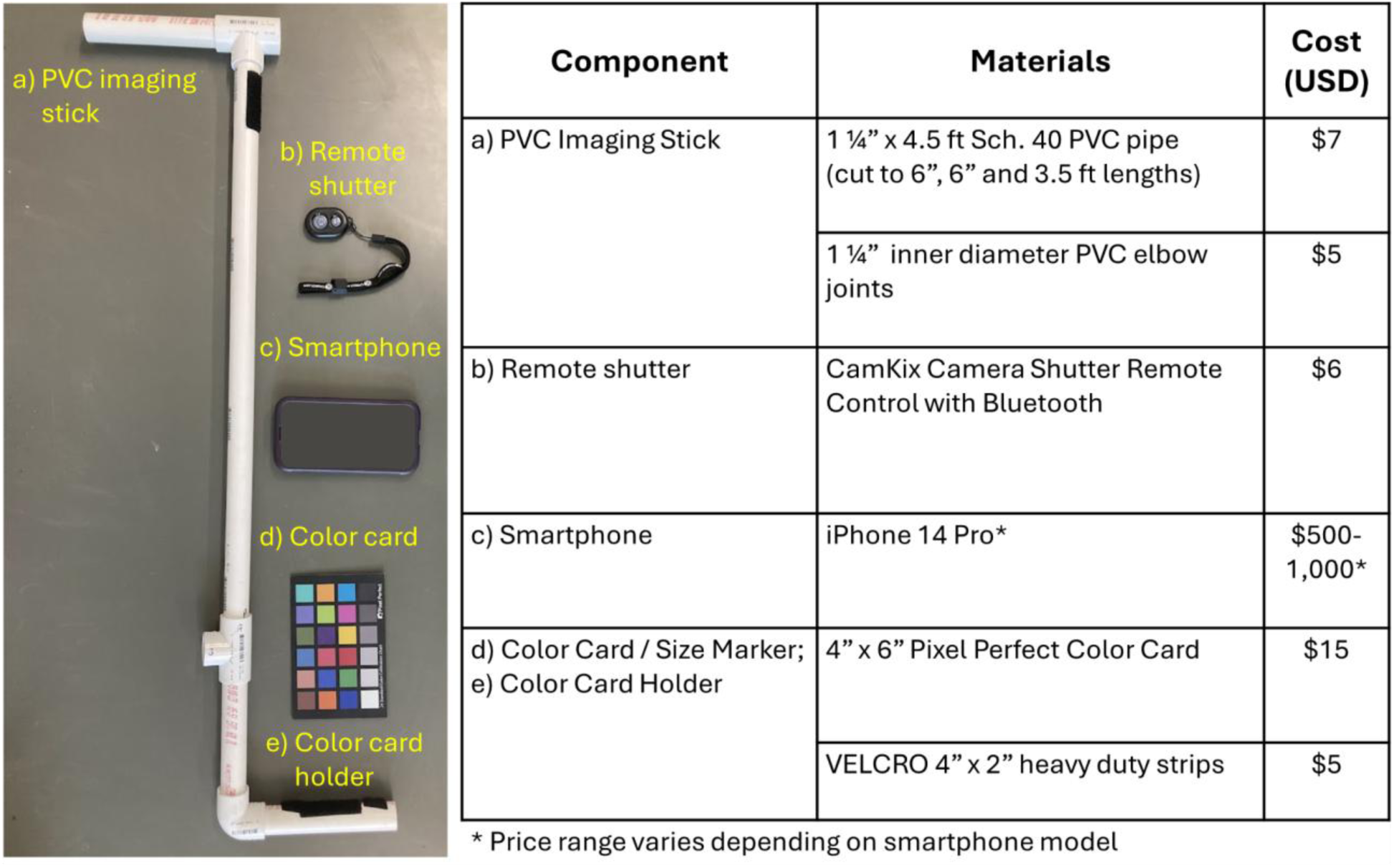
List of materials and their approximate cost for the smartphone image capture system (SICS).

**Figure 2.**
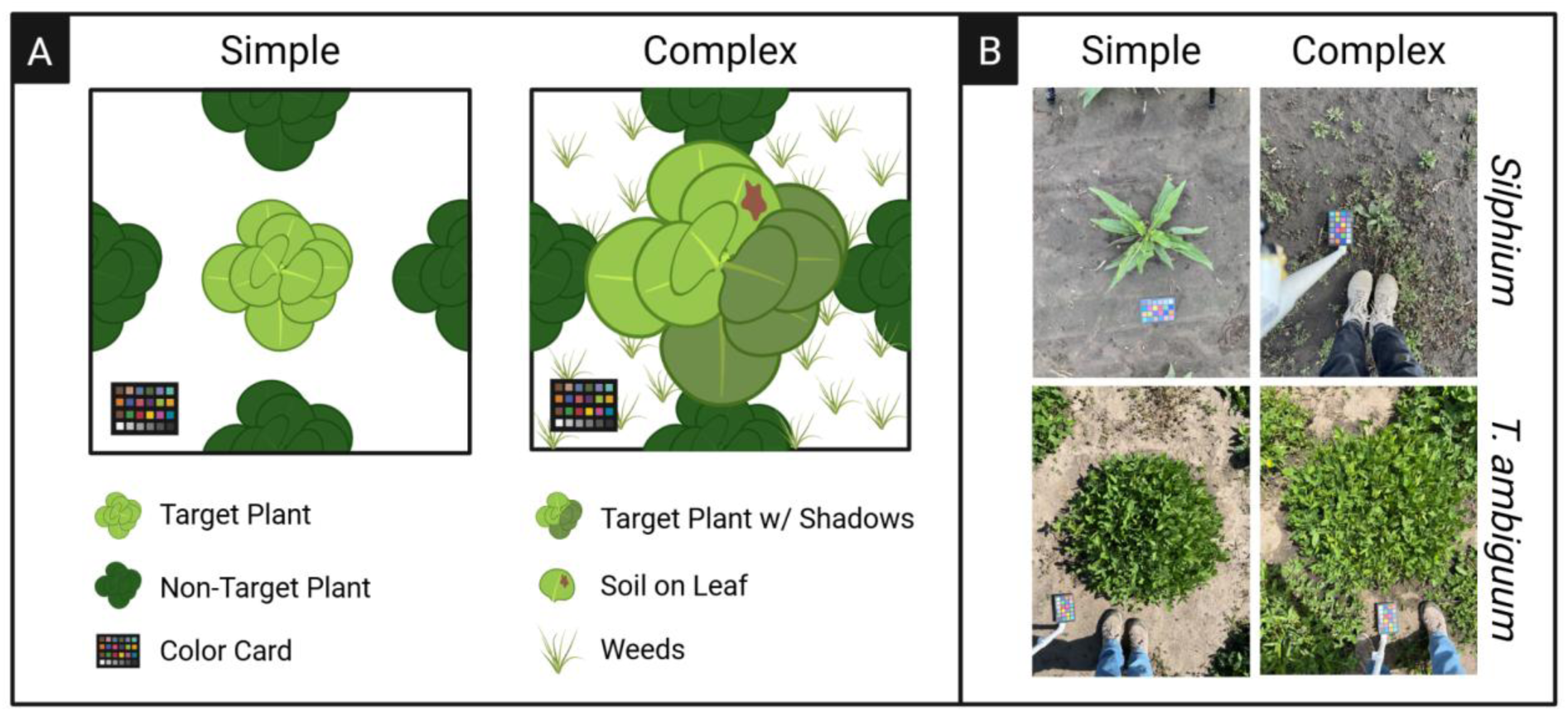
A) Illustration depicting simple and complex image compositions. Simple images were those characterized by little background noise (e.g., weeds, non-target plants, or organic debris), few shadows obscuring target plants, and no non-target plants touching target plants. Complex images were those characterized by any of the following conditions: i) high background noise from weeds, non-target plants, or organic debris, ii) shadows obscuring target plant, or iii) non-target plants touching target plants. All images contained a color card as a size reference. Illustration created using https://BioRender.com.B) Examples of simple and complex images collected from *Silphium* and *T. ambiguum* spaced plant nurseries.

### 2.3 Simple vs. Complex Image Test Sets

The spaced plant nursery layout, coupled with differences in shoot morphology between *T. ambiguum* and *Silphium*, allowed us to use the SICS to capture the variety of image conditions required to test the robustness of each image analysis pipeline to the range of image complexity encountered in the field (e.g., shoots occluded by shadows or soil debris, touching of target and non-target plants) (Fig 2). The variability in shoot height within *T. ambiguum* crowns (approx. 5 - 30 cm in height) made these plants more susceptible to having portions of the crown occluded by shadows in overhead images. Conversely, the prostrate growth of *Silphium* rosettes made them more susceptible to accumulating soil and debris on leaf tissue due to the proximity of the leaf to the soil surface. Further, the spaced plant layout of the *T. ambiguum* and *Silphium* nurseries allowed us to capture images where target plants were either touching or not touching non-target plants, such as neighboring plants or weeds.

Based on these conditions, images were classified manually as either ‘simple’ or ‘complex’ (Figure 2). Simple images were those with little background noise (e.g., weeds, non-target plants, or organic debris), few shadows obscuring target plants, and no non-target plants touching target plants. These images were more representative of controlled imaging conditions and were expected to pose fewer challenges for automated image analysis. Complex images were those characterized by any of the following conditions: i) high background noise from weeds, non-target plants, or organic debris, ii) shadows obscuring target plant, or iii) non-target plants touching target plants. The composition of complex images was more representative of the conditions encountered in field environments that were expected to hamper automation of downstream image analysis.

Two sets of images were used to evaluate the performance of image analysis pipelines. The first set was a subset of images representing each image composition type (i.e., simple or complex) and was used to assess the accuracy of each image analysis pipeline. For each species, a total of 50 simple and 50 complex images were randomly selected from the full image sets from each image collection year (Table 1), yielding a total of 100 images of *T. ambiguum* and 100 images of *Silphium* in this image set. The second image set contained images of four replicates each of three *T. ambiguum* genotypes imaged at two timepoints (March 27, 2025 and April 22, 2025) and tested whether mean plant area across biological replicates estimated using the PlantCV and Biodock AI pipelines would produce similar relative biological trends to those of ImageJ, regardless of image analysis pipeline accuracy. Images of *T. ambiguum* replicates collected on March 27, 2025, were plants with simple image compositions, and images collected on April 22, 2025, were plants with complex image compositions. In total, this image set contained 96 images (3 genotypes x 4 replicate individuals imaged per genotype x 2 image compositions).

### 2.4 Image Analysis Pipelines

We tested the ability of two image analysis pipelines to generate accurate and biologically meaningful plant area estimates from simple and complex images. The first pipeline was fully automated and used PlantCV, an open-source python-based image segmentation tool designed specifically for the extraction of plant traits from images (Gehan et al., 2017). A second, semi-automated pipeline was developed using Biodock AI, a commercially available AI-assisted image segmentation platform (Biodock AI, 2024) that enables manual correction of segmented objects. An ImageJ pipeline, which relied solely on manual image annotation, was used as a baseline to which the performance and output of the PlantCV and Biodock AI image analysis pipelines were compared (Elliot et al., 2022; Schindlein et al., 2012). We tracked the time between image collection and data availability in order to assess efficiency of the two pipelines.

Each pipeline was used to extract pixel counts from the target plant and color card chips, which are of known size so can be used to calculate actual plant area. Using PlantCV, a custom python script was written to quantify the target plant pixels in each image, which was then converted to actual plant area. Briefly, a binary mask was created using a dual channel threshold to isolate green plant pixels from background pixels. Green plant pixels were converted to white and background pixels were converted to black. A circular region of interest (ROI) was defined over the center of the image, approximately where the center of each target plant was located, and the binary mask was filtered to include only white pixels contained in or objects in the binary mask overlapping with the ROI. The number of white pixels contained in the filtered mask was extracted as the pixel count of the target plant. The pixel count of the color card was determined using the built-in detect color card function. Separate scripts were required to analyze each test image set for each image composition and plant species combination.

Similar to the PlantCV pipeline, the Biodock AI pipeline estimated plant area using the pixel area of the target plant and color card in each image. For each plant species, one SAM was trained to identify plants and color cards in each image. Following segmentation, each image was manually edited within the Biodock AI web-based platform to correct boundaries of the target plant and color card objects and eliminate non-target plant objects. Separate models were trained for each plant species. Within plant species, the same model was used to analyze simple and complex images. Analysis time was estimated as the time required to deploy the model and apply corrections to objects in images.

In ImageJ, the target plant and color card were manually traced using the freehand and polygon selection tools. Pixel counts were extracted from these selections using the Measure function.

Image analysis pipeline outputs were assessed using the plant area phenotype. The target plant and color card pixel counts generated from each image analysis pipeline were used to calculate the actual plant area (m^2^) of each target plant as follows:

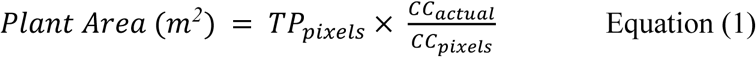

where TP_pixels_ and CC_pixels_ are the pixel counts of the target plant and color card, respectively, and CC_actual_ is the actual measured area of the color card (0.015 m^2^).

Image processing times were recorded as the time required to extract pixel counts of target plants and color cards from simple and complex images using each image analysis pipeline. For the PlantCV pipeline, the image processing time was the amount of time required to execute the custom PlantCV scripts to extract pixel counts for target plants and color cards. For the Biodock AI pipeline, the image processing time was the amount of time required to deploy the segmentation model and apply manual corrections to segmented target plants and color cards. For ImageJ, the image processing time was the amount of time required to manually extract pixel counts from target plants and color cards in each image.

### 2.5 Data Analysis

The accuracy of the PlantCV and Biodock AI pipelines compared to ImageJ was assessed using the first image set, which contained 50 simple and 50 complex images of each plant species. Accuracy was evaluated using simple linear regression to determine the *Pearson correlation coefficient* between the plant area estimated using ImageJ and that of the PlantCV or Biodock AI pipelines. Correlations were considered significant when *p* < 0.05. The mean relative difference between plant area estimated using ImageJ and the PlantCV or Biodock AI pipelines was also computed for each species and image composition combination.

It is well understood that automated and semi-automated image analysis methods are imperfect, and researchers ultimately aim to extract trait data from images that are of suitable accuracy for answering the biological questions of interest (Gehan and Kellogg, 2017).

Therefore, lower accuracy of image analysis pipelines may be acceptable, provided the image analysis pipeline retains relative trends in the plant trait of interest. To this end, we also tested whether mean plant area across biological replicates estimated using the PlantCV and Biodock AI pipelines would produce similar relative biological trends to those of ImageJ, regardless of image analysis pipeline accuracy. For this test, the second image set was used, which contained simple and complex images of individual *T. ambiguum* replicates for three genotypes. For each image composition, the effect of *T. ambiguum* genotype, image analysis pipeline and their interaction on mean plant area was tested using two-way analysis of variance (ANOVA). A significant interaction (*p* < 0.05) between *T. ambiguum* genotype and image analysis pipeline indicated trends in plant area between image analysis pipelines were significantly different.

All data analysis and figure creation was performed using R (version 4.3.1, R Core Team, 2024) using the *here* (ver. 1.0.1, Muller 2020), *tidyverse* (ver. 2.0.0, Wickham et al., 2019) *ggpubr* (ver. 0.6.0, Kassambara, 2025) and *ggplot2* (ver. 3.4.4, Wickham et al., 2016) packages.

## 3 Results

The smartphone image capture system (SICS) cost less than $1,050 (Figure 1), which included the smartphone, color card and materials for the PVC imaging stick. The SICS was used to collect over 12,000 images in spaced plant nurseries during the 2023-2025 growing seasons (Table 1). Using the SICS, image collection took just 6 to 24 minutes per 100 plants in the spaced plant populations depending on the walking speed of the photographer and the plot design, enabling the collection of images over a high temporal frequency (Table 1). The smartphone images were of high enough quality to be analyzed using both automated (PlantCV) and semi-automated (Biodock AI) image analysis pipelines.

The amount of time required to generate plant area data of target plants varied by image analysis pipeline (Table 1). Taking into account the human user time, the Biodock AI pipeline segmented target plant and color card pixels 22 and 36 times faster than manual image segmentation using ImageJ for *Silphium* and *T. ambiguum*, respectively. Similarly, the Biodock AI pipeline segmented target plant pixels from images 28 and 81 times faster than the PlantCV pipeline. However, it is important to note the analysis times associated with PlantCV were wholly computational, required no active time from a human user, and are dependent on the amount of compute resources allocated to the pipeline. If running PlantCV programs in parallel, analysis times can be greatly reduced. The Biodock AI pipeline required approximately 15 minutes of human user time per 100 images and analysis with the ImageJ pipeline was entirely performed by a human user.

Both the PlantCV and Biodock AI image analysis pipelines produced similar plant area estimates as ImageJ for simple images (Figure 3A, Figure 3B). For *T. ambiguum* and *Silphium* images, the correlation between the plant area estimated by ImageJ and either the PlantCV or Biodock AI pipelines was significant (p < 0.05), and the Pearson correlation coefficient was greater than 0.8. Together, these results indicate strong positive correlation between the ImageJ and alternative image analysis pipeline outputs. These results were supported by the low relative difference in plant area estimated using the ImageJ pipeline and PlantCV or Biodock AI pipelines. Simple *Silphium* images were on average within 11% and 7% of the ImageJ-generated plant area for the PlantCV and Biodock AI pipelines, respectively. For simple images of *T. ambiguum*, the relative difference between ImageJ and the PlantCV or Biodock AI pipelines increased slightly to 31% and 8%, respectively, but remained low.

**Figure 3.**
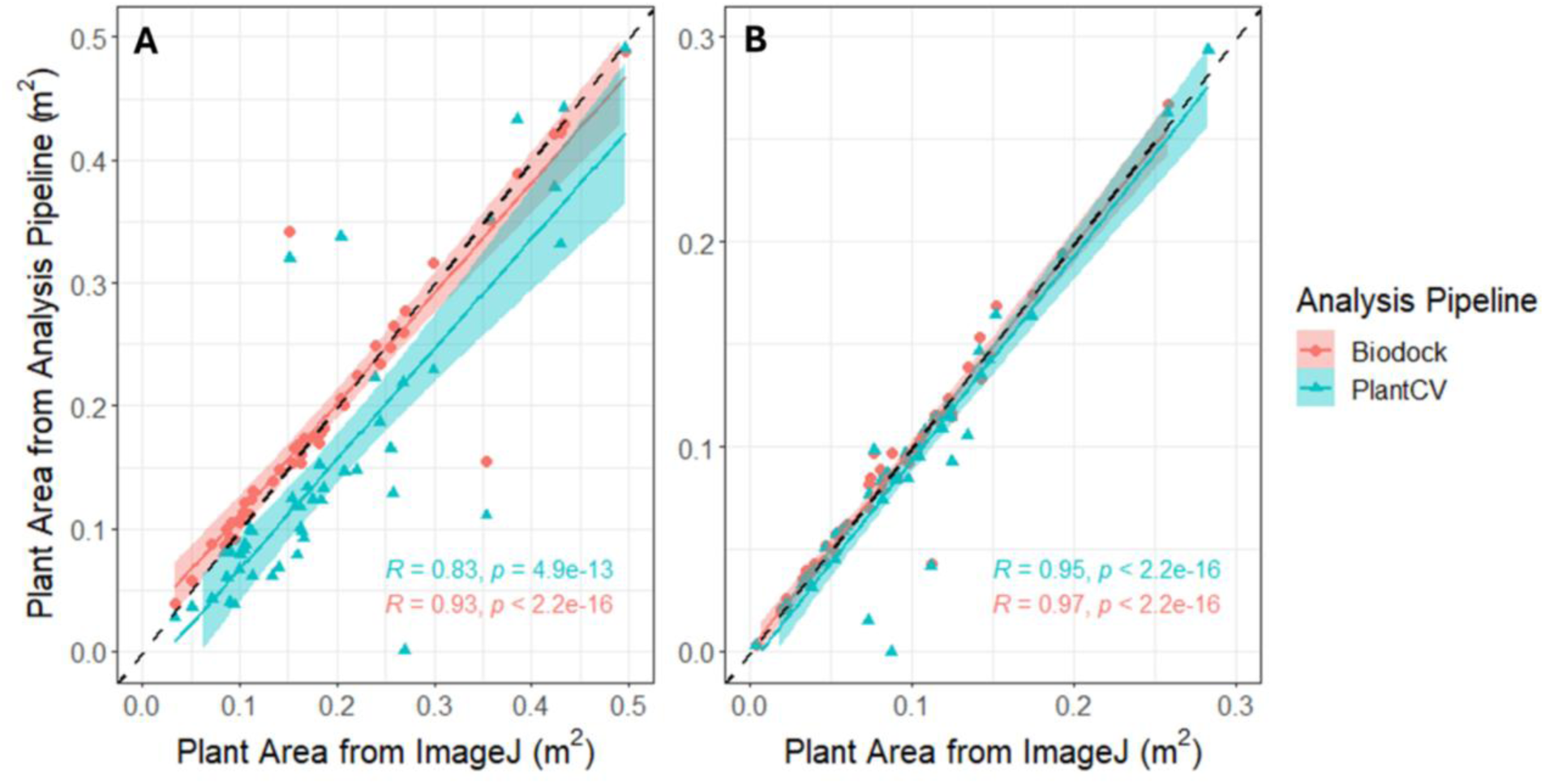
Pearson correlation coefficients and significance of linear regression between plant area (m^2^) estimated by ImageJ and PlantCV or Biodock AI pipelines for simple images of *T. ambiguum* (A) and *Silphium* (B) plants.

Compared to simple images, the accuracy of both PlantCV and Biodock AI pipelines declined with complex images. Though plant area estimated using PlantCV and Biodock pipelines remained significantly positively correlated to that of ImageJ (Figure 4A, Figure 4B), the Pearson correlation coefficients generally declined. The correlation between plant area estimated by ImageJ and the Biodock AI pipeline was similar for simple (R=0.93) and complex (R=0.96) images of *T. ambiguum*, this correlation declined from simple (R=0.83) to complex (R=0.59) images when using the PlantCV pipeline (Figure 4A). Similar trends were seen in the correlation between plant area estimated using ImageJ and PlantCV or Biodock AI pipelines for complex *Silphium* images, where Pearson correlation coefficients declined marginally when compared to simple images. In line with the decline in Pearson correlation coefficient, the relative differences between plant area estimated using ImageJ and the PlantCV or Biodock AI pipelines for complex images generally increased compared to simple images for both plant species (Table 2). This increase was particularly notable for complex images of *Silphium*, where the relative difference between plant area estimated using ImageJ and the PlantCV and Biodock pipelines was 41% and 93%, respectively, nearly 6 and 8 times greater than that for simple images.

**Figure 4.**
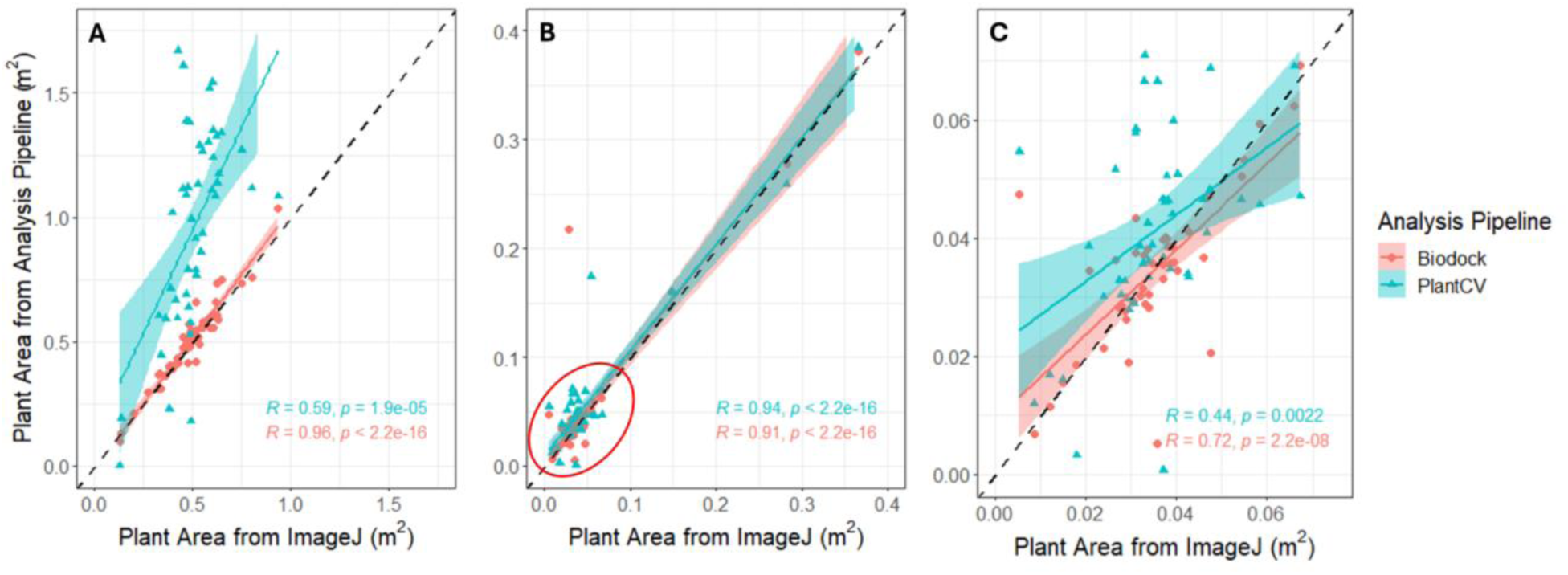
Pearson correlation coefficients and significance of linear regression between plant area (m^2^) estimated by ImageJ and PlantCV or Biodock AI pipelines for complex images of *T. ambiguum* (A) and *Silphium* (B,C) plants. Panel 4C shows the points contained in the red circle of panel 4B.

**Table 2.** Mean plant area (cm^2^; ± standard error) and relative difference (%) of plant area between ImageJ and PlantCV or Biodock AI pipelines.

|  | Simple |  | Complex |  |
| --- | --- | --- | --- | --- |
|  | Plant Area<br>(cm<sup>2</sup>) | Relative Difference<br>(%) | Plant Area<br>(cm<sup>2</sup>) | Relative Difference<br>(%) |
| *Silphium* |  |  |  |  |
| ImageJ | 992 ± 95 | 0 | 490 ± 86 | 0 |
| Biodock | 998 ± 99 | 7 | 527 ± 95 | 41 |
| PlantCV | 930 ± 100 | 11 | 687 ± 142 | 93 |
| *T. ambiguum* |  |  |  |  |
| ImageJ | 2145 ± 190 | 0 | 4904 ± 211 | 0 |
| Biodock | 2158 ± 182 | 8 | 5034 ± 228 | 7 |
| PlantCV | 1724 ± 201 | 31 | 9647 ± 590 | 95 |

Despite the differences in accuracy between ImageJ and the PlantCV and Biodock AI pipelines, particularly for those of complex images, trends in plant area across biological replicates remained consistent (Figure 5A, Figure 5B). For simple images (Figure 5A), the significant main effects of plant genotype (*p* = 0.012) and analysis pipeline (*p* = 0.018) on plant area were significant (*p* < 0.05), with PlantCV pipeline producing significantly lower plant area estimates than the ImageJ or Biodock AI pipelines. However, the interaction between analysis pipeline and plant genotype was not significant (*p* = 0.012). Similar results were observed for complex images, where neither the main effects of plant genotype (*p* = 0.338) and analysis pipeline (*p* = 0.270), nor their interaction (*p* = 0.942) were significant (Figure 5B). The nonsignificant interaction between plant genotype and analysis pipeline for both simple and complex images indicated that, although mean plant area estimates differed across image analysis pipelines, the image analysis pipeline did not significantly alter trends in plant area across biological replicates when compared to ImageJ.

**Figure 5.**
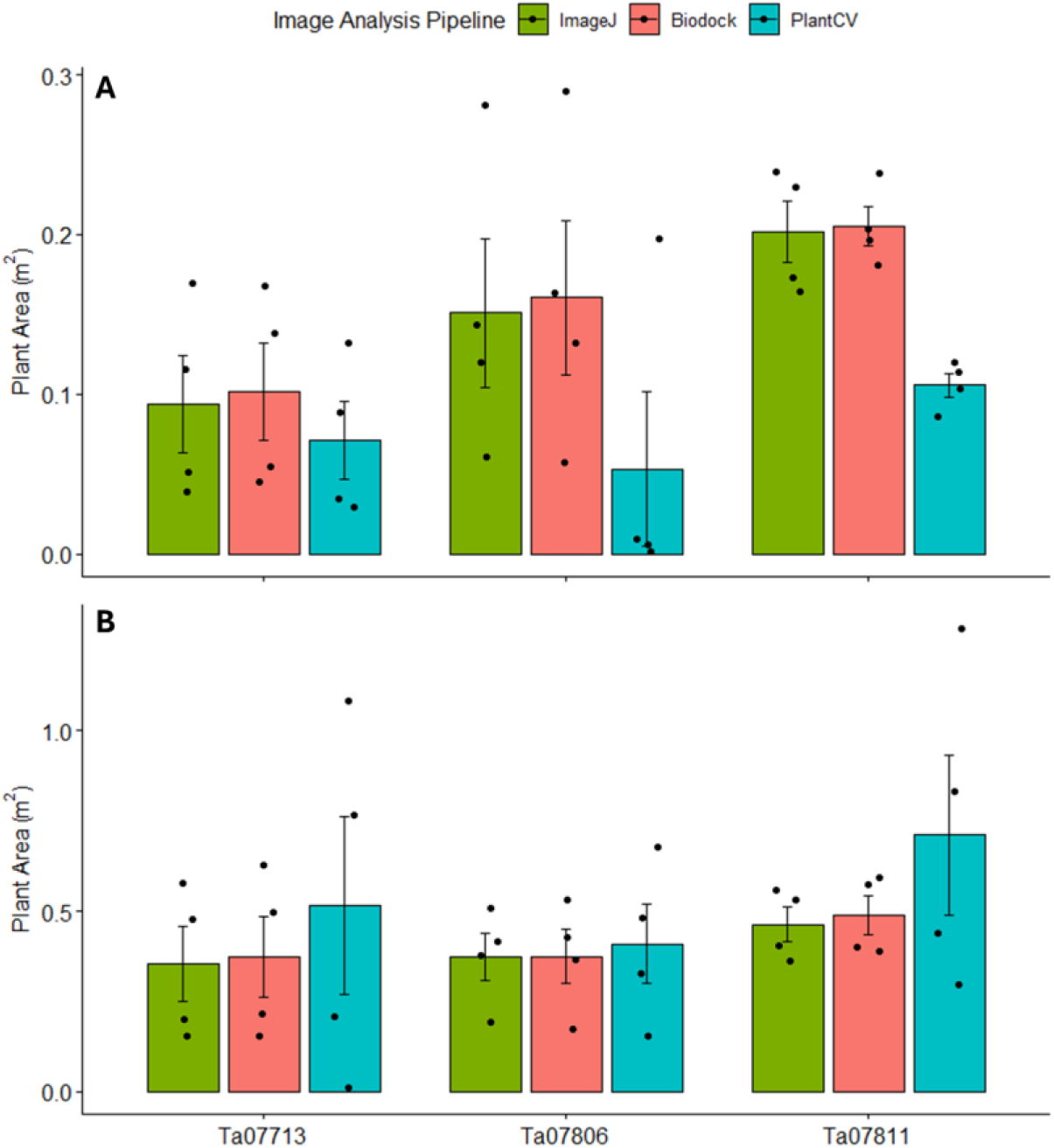
Mean (± standard error) plant area (m^2^) of three *T. ambiguum* genotypes for simple (A) and complex (B) images.

## 4 Discussion

Barriers associated with existing field image capture systems, such as unoccupied aerial vehicles (UAVs) and ground-sensing vehicles (GSVs), limit the adoption of image-based phenotyping in field experiments. For example, in order to operate UAVs, researchers must adhere to strict federal aviation policies and receive specialized training and licensure (Guo et al., 2021; Gano et al., 2024). Though GSVs are not subject to the same regulatory pressures as UAVs, GSV operation still requires specialized technical expertise for operation (e.g., tractors, utility vehicles and custom-built rovers) and mechanical or engineering expertise for design and construction (Jiang et al., 2018; Busemeyer et al., 2013). In contrast, the regulatory and technical barriers to using the SICS are low. Composed only of a smartphone, polyvinyl chloride (PVC) pipes, color card and remote shutter, the SICS relied on equipment that is easy to use, readily available, and requires no specialized training or licensure to operate. This design is expected to enable researchers to easily employ the SICS in a variety of field research settings, expanding the accessibility of field-based phenotyping to a wider range of researchers.

Due to the streamlined design of the image capture stick and ubiquitousness of smartphones, the SICS also ameliorated cost-related barriers to in-field image-based phenotyping. The cost of constructing the SICS was under $1,050 USD. When not accounting for the cost of the smartphone, which many researchers already have access to, this cost was reduced to less than $50 USD. Comparatively, the cost of UAVs alone ranges from $500 - $5,000 USD (Jang et al., 2020), with an additional tens of thousands of USD incurred by labor and training required to operate UAVs (Reynolds et al., 2019). Though not explicitly reported, the cost of SICS was also expected to be greatly reduced compared to plant- and plot-level GSVs. Even when using pre-owned tractors or utility vehicles, the assembly of GSVs often require purchase of cameras and mechanical infrastructure (e.g., imaging boxes, camera housing, etc.) that incur costs (Jiang et al., 2018; Busemeyer et al., 2013). The greatly reduced cost of SICS compared to existing in-field image capture systems lowers the barrier of entry for using image capture systems in field studies, enabling wider adoption of image-based phenotyping in plant breeding and research programs.

The SICS presented a low-barrier alternative to existing image capture systems, but its application may be limited by the size of the field experiment. While the SICS enabled the collection of over 10,000 images each field season, image capture in this study only occurred in spaced plant nurseries, where individual target plants were less than 1 m apart. When imaging experiments with a large spatial footprint, the SICS may be less practical and expedient than UAVs and GSVs, which can collect images of acres of land over a short duration. While UAVs and GSVs may still be advantageous over the SICS when imaging experiments that cover large areas, the SICS is still expected to present a readily accessible and expedient alternative to existing field image capture systems in studies of individual plants and small spaced-plant nurseries. Beyond constraints of scale, we recognize that SICS may not be amenable to all planting designs where individual organisms are not readily discernible (plots and mixed plantings), or for those including taller plants with complex above-ground architectures. Further, though the SICS is low-cost and easy-to-use, the SICS is still a manual imaging process that requires an operator to collect images, which can be associated with costs for human hours, as well.

Following image collection in the field, low turnaround times between image collection and plant trait data availability presents an additional barrier to adopting image-based phenotyping methods in field studies. Lacking widely available user-friendly image analysis pipelines, processing UAV images is time consuming and computationally intensive (Gano et al., 2024; Bongomin et al., 2024), sometimes requiring months of human labor to develop analysis pipelines capable of extracting plant trait information. Even with ground-level imagery, field images are often characterized by conditions (e.g., target plants touching non-target plants, shadows and soil obscuring leaf tissue) that challenge the use of high-throughput image analysis pipelines for plant trait extraction. Some have attempted to reduce image processing times by altering field imaging methods, with adaptations ranging from imaging plants against a black backboard to attaching mobile lightboxes to image capture systems (Dang et al., 2023; Busemeyer et al., 2013; Jiang et al., 2018). However, such adaptations add additional complexity to field image collection methods and ultimately optimize image processing times at the expense of expedient field image collection.

Instead of modifying field imaging methods to reduce image analysis times, this study demonstrated the use of two image analysis pipelines to rapidly produce plant trait information from a variety of field image conditions. Using the PlantCV and Biodock AI pipelines, image analysis times were reduced from weeks or months to minutes or hours, with much of the reported processing time being entirely automated and without the need for human labor.

Further, both image analysis pipelines relied on readily available image segmentation tools, easing barriers associated with limited availability of UAV and GSV image processing software (Gano et al., 2024). Coupled with the low image collection times, the image analysis pipelines demonstrated in this study present a rapid and accessible option to expedite field phenotyping for individual plant or small-plot research studies.

Importantly, reductions in image collection and processing times did not come at the expense of plant trait data accuracy. When analyzing simple images, both the PlantCV and Biodock AI pipelines produced accurate plant area estimates. As PlantCV and Biodock AI were primarily designed to extract features from images collected under controlled conditions, these results were in line with expectations. However, managing field sites to maintain simple image compositions is rarely feasible for larger scale experiments. More often, images collected in the field are characterized by complex conditions that complicate automated and semi-automated image analysis, such as target plants touching non-target plants and occlusion of leaf material by shadows or soil. When analyzing complex images, the accuracy of the image analysis pipelines tended to decrease, particularly for the PlantCV pipeline where trends suggested the PlantCV pipeline overestimated plant area. These results were likely due to the thresholding and object selection tools used to isolate target plants, in combination with the greater proportion of target plants touching non-target plants. Using PlantCV, the target plants were isolated by first selecting green pixels, then selecting continuous areas of green pixels that intersected a region of interest outlined over the center of the target plant. With the complex images featuring target plants touching non-target plants, thereby creating a continuous line of green pixels, this segmentation method resulted in non-target plants being identified as target plants, artificially inflating the plant size estimates from these images. These results suggested alternative image segmentation methods within PlantCV, or other pre-processing, may increase accuracy of phenotype extraction.

Despite the observed decline in accuracy, a key finding from this study was that useful plant trait data were extracted from images using these pipelines even when analyzing complex images. The lack of significant difference in relative trends across biological groups (i.e., genotype) suggests that either pipeline could be used to extract plant trait information regardless of accuracy. However, although both pipelines were suitable for analyzing both simple and complex images in this study, researchers interested in employing these methods on complex images should evaluate the performance of these pipelines for their specific research needs prior to employing these methods on a larger scale.

While both image analysis methods employed in this study produced accurate biologically relevant trait measurements, they each have tradeoffs. Biodock AI (and other proprietary AI segmentation platforms) incurs a cost related to computational resources and the resulting models are often not easily reused, making incremental updates or foundational models challenging. Open-source software like PlantCV is often programmatic and users need to supply the computation resources, creating a barrier to use. Based on the results presented here, future improvement could involve combining easy to use model training platforms like Biodock AI with the flexibility and downstream breadth of analysis modules already implemented in PlantCV. Binary masks resulting from machine learning image segmentation methods can already be directly used by PlantCV, but automation of this has not been optimized.

Additionally, PlantCV already has functions to train two types of machine learning classifiers for image segmentation: naive bayes (supervised) and k-means clustering (unsupervised), both of which are computationally lighter than models like U-Net and Segment Anything. Because PlantCV is python-based and implemented typically through either Jupyter notebooks or command-line parallel job submission, there is no intrinsic barrier to incorporating more complex models for segmentation upstream of trait extraction. Such an update could provide further accuracy for traits like size and allow for extraction of the more complex morphological traits that PlantCV is already capable of analyzing.

## 5 Conclusion

Numerous financial, technical and computational barriers prevent the widespread adoption of image-based field plant phenotyping methods, particularly for small research groups or individuals. In this study, we presented an inexpensive and user-friendly smartphone image capture system (SICS) that enabled the rapid collection of plant images from three field experiments. Though computational bottlenecks can prolong the time between image collection and trait extraction, this study further demonstrated the use of two easily accessible image analysis pipelines to expediently extract accurate plant trait information from images. It is expected that the image capture and analysis methods detailed in this study could be easily adopted by research groups to alleviate the phenotyping bottleneck. This is especially important for plant breeding and crop development programs, including those focused on emerging herbaceous perennial crops. Such methods will better enable phenotypes to be collected on a timeline that enables rapid advancement of crop germplasm to meet current economic and environmental challenges.

## Acknowledgements

This work was funded through the New Roots for Restoration Biology Integration Institute (NSF 2120153). We thank the Danforth Center Field Research Site field crew (Terry Beeler, Nelson Curran, Kevin Hava) as well as Danforth Center researchers Shayla Gunn, Finn Reeves, Tyler Thrash, Sravani Valligari, Isabella Vergara, Tuany Volz, and the Danforth Center Data Science facility (RRID:SCR_027573) for their assistance with the field preparation/transplanting, field maintenance, image collection and image processing that supported this work.

